# Mechanism of Nanomaterial-Induced Lipid Droplet Formation in *Raphidocelis subcapitata* is Mediated by Charge Properties

**DOI:** 10.1101/2025.02.11.637709

**Authors:** Emma McKeel, Hye-In Kim, Su-ji Jeon, Britta McKinnon, Juan Pablo Giraldo, Rebecca Klaper

## Abstract

Increasing the production of renewable energy will be critical to achieving global sustainability goals in the coming decades. Biofuels derived from microalgae have great potential to contribute to this production. However, cultivating algae with sufficient neutral lipid content, while maintaining high growth rates, is a continual challenge in making algal-derived biofuels a reality. Previous work has shown that exposure to polymer-functionalized carbon dots can increase the lipid content of the microalgae *Raphidocelis subcapitata*. This study investigates this finding, aiming to determine the mechanisms underlying this effect and if altering nanoparticle surface charge mediates the mechanism of action of the carbon dots used. Carbon dots with both negative and positive surface charges were added to microalgal cultures, and the impacts of this exposure were analyzed using high-content imaging, growth measurements, and chlorophyll content measurements. Results indicate that positively charged carbon dots induce a nano-specific increase in lipid content but also cause decreases in growth. Additionally, the mechanism of action of each nanoparticle was examined by conducting a morphological comparison to known treatments. This analysis showed that negatively charged carbon dots cause similar impacts to *R. subcapitata* as nitrogen deprivation, a known mechanism by which lipid droplet content can be increased. The findings of this study suggest that carbon dots may have surface charge dependent effects on the lipid metabolism of *R. subcapitata*. Future work should consider the use of carbon dots with varied surface charge densities for enhancing algae biofuel production in bioreactors.

## Introduction

Reducing the global risks posed by climate change requires immediate changes, especially to our energy infrastructure. Luckily, many renewable fuel source options could be exploited in coming years. Microalgae are one source of renewable fuel which has been increasingly studied for this purpose. The U.S. Department of Energy estimates that the U.S. has the potential to produce 191 million tons of microalgal dry weight per year. Furthermore, algae do not require arable land for production and can be grown in waste or saline water [1]. These factors suggest great potential for microalgal derived biofuels to contribute to global production of sustainable energy. Despite this promise, many challenges remain on the path to making algal biofuels economically feasible. Biofuels derived from algae are produced by extraction of neutral lipids, especially triacylglycerols (TAGs) from the cell. These lipids are preferred sources of biofuels due to their high percentage of fatty acids [2]. However, TAGs are accumulated to the highest extent under stress conditions, such as nutrient deprivation or photo-oxidative stress [3]. This presents a challenge for cultivating algae for biofuel production: how can TAG content be increased without inducing too great of stress levels, which could inhibit cell growth?

Nanoparticles, defined as a particle with a diameter between 1 and 100 nm, have been primarily studied for their toxic effects on microalgae. In recent years, though, the use of nanoparticles in algal biofuels research has grown. Tested applications of nanoparticles in algal-based biofuels research include the addition of iron and magnesium sulfate nanoparticles to increase lipid production [4, 5] and downstream applications of nanoparticles for more efficient lipid harvesting and removal [6]. Another nanoparticle which has been shown to increase microalgal lipid production is carbon dots (CDs). Carbon dots are a, organic, carbon-based (also containing hydrogen, oxygen, and nitrogen), versatile nanoparticle, with tunable surface chemistry and intrinsic fluorescence characteristics [7]. In addition to their use in imaging, carbon dots are increasingly being investigated for use in nano-enabled agriculture to enhance photosynthesis and serve as vessels for gene delivery to chloroplasts [8, 9] and targeted chemical delivery to the plant vasculature mediated by biorecognition [10]. A 2024 study of three polymer functionalized carbon dots: Polyethylenimine (PEI), carboxylated polyethylenimine (CP) and polyvinylpyrrolidone (PVP) CDs, found that carbon dots also have the potential to increase lipid content in the green microalgae *Raphidocelis subcapitata* [11]. Positively charged PEI-CDs increased lipid droplet content at a concentration of 10 ug/L, while reducing growth by only 11.69 ± 4.50%. Exposure to 10 or 50 ug/L of negative CP-CDs or 50 ug/L of near neutral PVP-CDs resulted in significant increases to lipid content without impacting growth. However, the mechanisms by which these impacts occurred remain unclear. This study seeks to further investigate the mechanisms underlying the impact of PEI and CP-CDs on lipid content in *R. subcapitata*. Although understudied in the field, *R. subcapitata* has recently been suggested as a species with great potential for biofuels production [12]. In fact, light intensity experiments comparing lipid content of *R. subcapitata* to that of *Chlorella vulgaris*, a popular candidate for biofuels production, found higher levels of lipids in *R. subcapitata* than in *C. vulgaris* [13]. Therefore, use of *R. subcapitata* in this study will provide further, much needed evidence regarding the species’ capacity for neutral lipid droplet production. It will also allow for direct comparison to previous results in which functionalized carbon dots impacted lipid droplet content in the species.

Carbon dots likely impact algal lipid metabolism via mechanisms specific to their surface charge and functionalization. Negatively charged CP-CDs have the potential to interact electrostatically with cations present in OECD 201 media. One of the most abundant of these is ammonium, the nitrogen source in OECD algal media [14]. Ammonium chloride is present at a final concentration of 15 mg/L in OECD media. Nitrogen deprivation is a well-documented method by which LD content can be increased in microalgae [3, 15–17]. Mechanistically, nitrogen deprivation remodels microalgal carbon metabolism, redirecting carbon towards lipid production [18] and impacting expression of acyltransferases [14]. It is hypothesized that electrostatic interactions between CP-CDs and the ammonium cations present in OECD media are the cause of increased LD content in CP-CD exposed microalgae. Binding of ammonium to CP-CDs is thought to reduce nitrogen availability to *R. subcapitata*, resulting in nitrogen deprivation and morphological effects consistent with it. To test this hypothesis, *R. subcapitata* cultures grown under low nitrogen conditions were morphologically compared to those exposed to CP-CDs. Random forest classification was utilized to compare CP-CD exposed cells to nitrogen deprived, control, and CP polymer exposed cultures. Classification was used to assist in determining which of these “known” treatments resulted in a morphology most similar to CP-CD exposure.

Consistent with the hypothesis that the mechanisms by which CDs increase LD content in *R subapitata* are determined by surface charge, we hypothesize that PEI-CDs impact LD content via a different mechanism than CP-CDs. This is supported by the toxicity observed in response to PEI-CDs, which did not occur after CP-CD exposure [11]. Additionally, PEI-CDs induced significant aggregation of *R. subcapitata* cells, another charge-specific response not observed with CP-CDs. The presence of cell aggregation indicates the potential for a shading effect to be at play. As cells aggregate with PEI-CDs and each other, light availability to individual cells could be decreased, causing growth inhibition and other negative effects. Shading has been previously documented as a mechanism of action by which nanoparticles, such as graphene oxide, impact microalgal growth [19]. It is less clear if nanoparticle-related shading could impact LD content. To first determine if PEI-CDs potentially impact *R. subcapitata* via shading, chlorophyll-a content was tested. If shading is occurring, a compensatory increase in chlorophyll-a should be observed, as is seen when algae are grown in low light conditions [20]. Chlorophyll-a content was compared among cells grown under control conditions, exposed to PEI-CDs or PEI polymer, and grown under reduced light conditions. Further, a morphological comparison was conducted among these treatments to determine if PEI-CDs induce a morphological response similar to growth under low light or exposure to the PEI polymer alone. Morphological profiling also allows for comparison between PEI and CP-CD exposed cells to identify differences in their impacts on *R. subcapitata*. These experiments provide additional information on the mechanisms by which PEI-CDs impact microalgae and thereby their LD content.

Previous studies found morphological changes in response to CD exposure using high-content imaging (HCI). Here, we expand these findings by comparing CD exposure to treatments with known mechanisms of action. High-content imaging is briefly defined as the use of automated, high-throughput fluorescence imaging and image analysis to identify phenotypic changes, especially in a cell culture. Having emerged from medical research, HCI is now gaining popularity in environmental research, including in microalgae [11, 21, 22]. In this work, images obtained via HCI imaging will be examined using two image analysis programs followed by a simple statistical analysis to determine how *R. subcapitata’s* lipid content is impacted by exposure to carbon dots or known treatments. The use of HCI will therefore not only provide useful data but will also serve as a demonstration of the use of this technology in microalgal studies.

## Methods

### 2.1 Starter Strain and Culture

The microalgae *Raphidocelis subcapitata* (also known as *Selenastrum capricornutum*) was obtained from the UTEX Culture Collection of Algae (UTEX 1648). *R. subcapitata* was cultured in OECD media [14] in an incubator at 24°C. Constant illumination at approximately 80 µmol s^-1^ m^-2^ was maintained. Flasks were kept on an orbital shaker at approximately 100 rpm. New cultures were inoculated weekly by adding 4 mL of algae to 400 mL of OECD media in a 1 L Erlenmeyer flask. Cultures were allowed to grow for four days prior to use in experiment setup.

### 2.2 Particle and Polymer Synthesis

Nanoparticles were synthesized as previously reported by Kim et al and McKeel et al [11, 23]. Briefly, the core carbon dot was synthesized by dissolving 1.92 g of citric acid and 2.40 g of urea in 2 mL of ultrapure water. Then, 1.35 mL of ammonium hydroxide (NH_3_·H_2_O, 30–33%) was added to the mixture with vigorous stirring for a few minutes until the ammonium hydroxide was fully dissolved. The reaction mixture was then transferred to a convection oven and heated at 180 °C for 1.5 hours. The mixture was allowed to cool to room temperature, then redissolved in 40 mL of ultrapure water while stirring at 150 rpm. Next, the solution was bath-sonicated at 37 Hz (60% power, Elmasonic P) for 30 minutes. This improved the dispersion of the nanoparticles. To remove any large aggregates, the solution was then centrifuged at 4500 rpm for 30 minutes, and the supernatant decanted. A dialysis membrane (molecular weight cutoff 3k, Spectrum) was used to further purify the CDs. Finally, the core CDs were transferred to a conical tube and once again bath sonicated for 30 minutes at 37 Hz, before being removed using a syringe filter (pore size 20 nm, Whatman). Core CDs were stored in a refrigerator after synthesis.

Core CDs were polymer functionalized as described by Kim et al and McKeel et al [11, 23]. To synthesize PEI-CDs, CDs were functionalized with PEI10k. To begin, 16 mL of 5 mg/mL core CD solution was suspended in a 20 mL glass vial. Using a NaOH solution (6 M) pH was modified to 12. To the solution, 3.2 mL of PEI10k (0.1 g/mL) was added while stirring vigorously. The mixture was transferred to an oven at 85 °C for 16 hours. Then, purification by extraction was completed. The mixture was then cooled to room temperature and transferred to a 500 mL Erlenmeyer flask. While stirring, an equal volume of ethanol and chloroform were added to the mixture, which was then vigorously mixed with a Voltex for 5 to 10 seconds and centrifuged at 4500 rpm for 5 minutes. After discarding the organic solvent layer, these extraction steps were completed 5 times. To remove unreacted polymer, 30k centrifugal filters were used 3 times.

Carboxylated PEI-CDs (CP-CDs) were synthesized by modifying the PEI-CD surface. To do so, 1.5 g of succinic anhydride was dissolved in 10 mL of DMF. The mixture was added to 30 mL of PEI-CD solution (1 mg/mL) in a 100 mL beaker. After stirring this reaction mixture for 16 hours, it was purified using the ethanol/chloroform extraction method described above, followed by filtration as described for synthesis of PEI-CDs.

The CP polymer was synthesized following a similar procedure to that used for CP-CDs. However, instead of using PEI-CDs, only PEI 10k polymer dissolved in DI water was used. The PEI solution was reacted with succinic anhydride under stirring for 16 hours. After the reaction, the mixture was purified using centrifugal filters (MWCO 10k) to remove the excess succinic anhydride and DMF.

### 2.3 Particle Characterization

Particle size and zeta potential were measured in TES buffer (10 mM, pH 7) using a Zetasizer (Malvern Panalytical) immediately after synthesis. Hydrodynamic diameter by volume (nm) and zeta potential (mV) were reported.

### 2.4 Experiment Setup

To ensure consistency in cellular concentration across replicates, chlorophyll fluorescence was used to estimate concentration prior to experiment setup. A standard curve was prepared for use in approximation of concentration in cells per milliliter. To do so, six samples of *R. subcapitata* were randomly diluted. 300 μL aliquots of each sample were added to a 96-well glass-bottom plate. The fluorescence of each of these samples was read using an Agilent BioTek Synergy H4 Hybrid Microplate Reader (Agilent Technologies, California, USA) at an excitation/emission of 440/670 nm. Fluorescence was corrected to a media control. The cell count in each random sample was then obtained using a hemacytometer and graphed opposite corrected fluorescence. This resulted in a linear equation relating chlorophyll fluorescence of a 300 μL sample to approximate cell count:

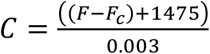

where C is cellular concentration in cells/mL, F is the fluorescence of the cell sample and F_c_ is the fluorescence of a cell-free media or media and nanoparticle control. This equation was used to estimate the cellular concentration of the stock from a 300 μL aliquot prior to exposure setup. Samples were normalized to a media-only control. Algae were then diluted to a final concentration of approximately 5×10^4^ cells/mL (actual starting concentrations were determined to range between 5×10^4^ and 5.62×10^4^ cells/mL).

To test the impact of nitrogen deprivation on *R. subcapitata*, four nitrogen-deplete OECD medias were prepared. In OECD media preparation, the nitrogen source, ammonium chloride, is added along with other nutrients as part of a stock solution. Three low-nitrogen vitamin stocks were prepared by reducing the mass of ammonium chloride added to 10, 25, or 50% of that required by the OECD media recipe. These stocks were used to make fresh nitrogen-deplete media each week with ammonium chloride concentrations of approximately 0.14 mM (50% N), 0.07 mM (25% N), and 0.028 mM (10% N). No other nutrient concentrations were modified. Experiments took place in 250 mL Erlenmeyer flasks at a final volume of 40 mL in the same conditions as the starter culture. Cell-free controls of each media type were prepared at experiment start and maintained in the same incubator as the experiment flasks.

CP-CD and PEI-CD exposures were carried out under identical conditions to low-nitrogen experiments. At exposure start, PEI-CDs were added to each exposure flask at concentrations of 0, 0.1, 1, 5, and 10 ug/L. Exposure concentrations were determined from toxicity data provided by McKeel et al [11], who showed that PEI-CD concentrations of 10 ug/L are sufficient to induce increased lipid content in *R. subcapitata.* Above this concentration, changes to lipid content were not observed. CP-CDs were added to exposure flasks in concentrations of 0, 5, 10, 25, and 50 ug/L. Similarly, these concentrations were derived from data published by McKeel et al [11], which demonstrated that the maximum CP-CD concentration inducing increased lipid content was 50 ug/L. At exposure start, a cell-free control for each CD dose was prepared. Each cell-free control had a final volume of 10 mL in a 15 mL conical tube. Cell-free controls were maintained in the same incubator as the exposure flasks.

Polymer-only controls were prepared in the same manner as CD exposures. Only the two highest doses tested for each CD were used for polymer-only controls. Cells were grown under the same conditions used for CD and low nitrogen experiments.

Low light experiments were carried out by placing a shade underneath the incubator light source, lowering the approximate photon flux reaching the algal flasks from 80 µmol s^-1^ m^-2^ to 10 µmol s^-1^ m^-2^. Cells were then grown in the same conditions as described for low nitrogen, CD, and polymer-only experiments.

### 2.5 Growth Inhibition

Cellular concentration was measured by removing a 300 μL aliquot from each experiment flask and calculating approximate cellular concentration using chlorophyll fluorescence (as described in *Experiment Setup*). These values were used to calculate percent growth inhibition as described by OECD Guideline 201 [13]. First, average specific growth rate was calculated using the equation:

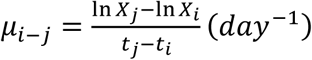

where μ*_i_*_-*j*_ is the average specific growth rate from time i to time j, *X_i_* is the biomass at time *i*, and *X_j_* is the biomass at time *j*.

Percent growth inhibition at 72 hours could then be calculated using the equation:

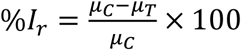

wherein %*I_r_* is the percent inhibition in average specific growth rate, μ_C_ is the mean average specific growth rate of control replicates, and μ_T_ is the specific growth rate for the treatment replicate. Control replicates were performed in conjunction with every experiment except those carried out in low light, as a full-light control was not possible in this setup. Any control replicates which did not increase in concentration by a factor of 16, an OECD guideline requirement, were removed. Thirteen weeks of 72-hour exposures were carried out in total, excluding one week of low light exposure. For final calculation of growth inhibition, one control was randomly selected from each of these weeks to include in analysis. This was done to account for any potential week-to-week variations in growth rate or culture conditions.

### 2.6 Chlorophyll-a Quantification

Chlorophyll-a content was measured for PEI-CD treated cells, in addition to those grown in low light conditions and in the presence of PEI polymer. PEI-CDs were targeted specifically for this assay due to the known toxicity of these particles and their documented aggregation effects.

Increased chlorophyll-a could indicate that aggregation is a major mode of toxicity, causing shading and a subsequent increase in chlorophyll. After 72 hours of CD exposure, algae was filtered from the exposure media using a GF/C grade glass microfiber filter (Whatman, United Kingdom). Filters were ground in 5 mL of extraction fluid (consisting of 68% methanol, 27% acetone, and 5% de-ionized water), before being transferred with a funnel to a 15 mL centrifuge tube. The used grinding tube, funnel, and pestle was rinsed with 5 mL of extraction fluid into the centrifuge tube to ensure complete collection of sample. Samples were then wrapped in aluminum foil and frozen for 24 hours at –20 °C. After 24 hours, samples were centrifuged for 5 minutes at 3000 to 4000 rpm. Without disturbing the sediment settled at the bottom of the centrifuge tube, a 5 mL pipette was used to fill a fluorometer tube just above halfway.

Fluorescence was then recorded. Two drops of dilute HCl were added to the sample, then fluorescence was recorded again. Chlorophyll-a content was then calculated using the equation:

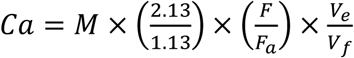

where C_a_ is chlorophyll-a content in μg/L, M is a multiplication factor determined by the scale ranged used on the fluorometer, F is the fluorescence before acid addition, F_a_ is fluorescence after acid addition, V_e_ is the volume of extract, and V_f_ is the volume which was filtered. Finally, chlorophyll-a was normalized by cell count for a final value in ng/cell.

### 2.7 Cell Staining and High-Content Imaging

At experiment end, 500 μL aliquots were removed from each flask for staining and high-content imaging. In a microcentrifuge tube, lipid droplets were stained with BODIPY 505/515 at a final concentration of 5 μM [24]. Nuclei were stained using NucBlue Live Ready Probes Reagent (Invitrogen, Massachusetts, USA). Cells were then incubated in the dark for 15 minutes. After staining, 300 μL aliquots were taken from each sample and dispensed into a 96-well glass-bottom plate for imaging. High-content image acquisition was completed using an ImageXpress Micro XLS Widefield High-Content Imaging System (Molecular Devices, California, USA), with images being obtained in the Cy5, GFP, and DAPI channels (Figure 1). In the Cy5 channel, the natural fluorescence of the chloroplasts was imaged. Stained nuclei were imaged in the DAPI channel, and stained lipid droplets in the GFP channel. Sixteen sites were imaged per well at 60x magnification. Twelve replicates of high-content imaging were carried out for each treatment.

**Figure 1.**
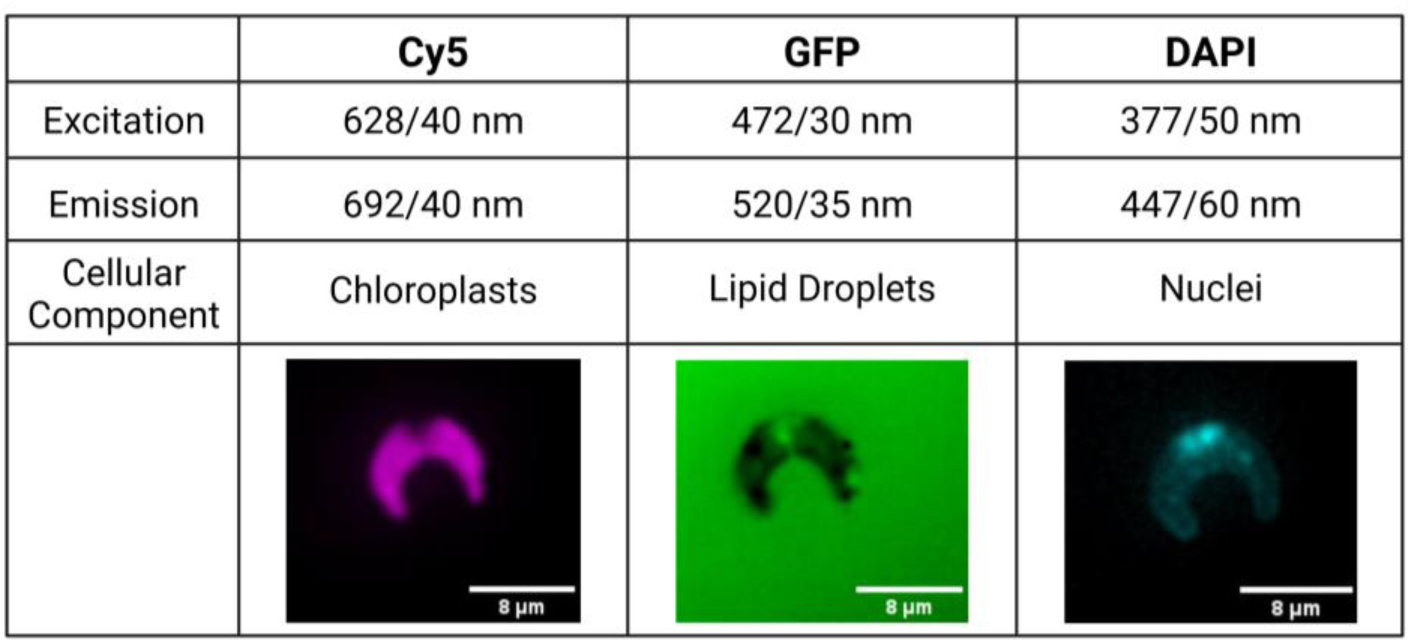
Example images accompanied by excitation and emission values of each fluorescence channel used in this study. Cell images edited in Fiji [40].

### 2.8 Image Analysis

Image analysis was carried out using both MetaXpress high-content image acquisition and analysis software (Molecular Devices, California, USA) and CellProfiler 4 [25]. Nuclei, lipid droplets, and chloroplasts were quantified using these techniques. Lipid droplet count data, normalized by cell count, was taken from MetaXpress, while all other measurements used in analysis were obtained from a CellProfiler pipeline. Additional documentation on these analysis pipelines is available in the Supplemental Information. Data was collected both per image and per object. For statistical review, per object data was averaged across each image and, subsequently, each replicate (Figure 2).

**Figure 2.**
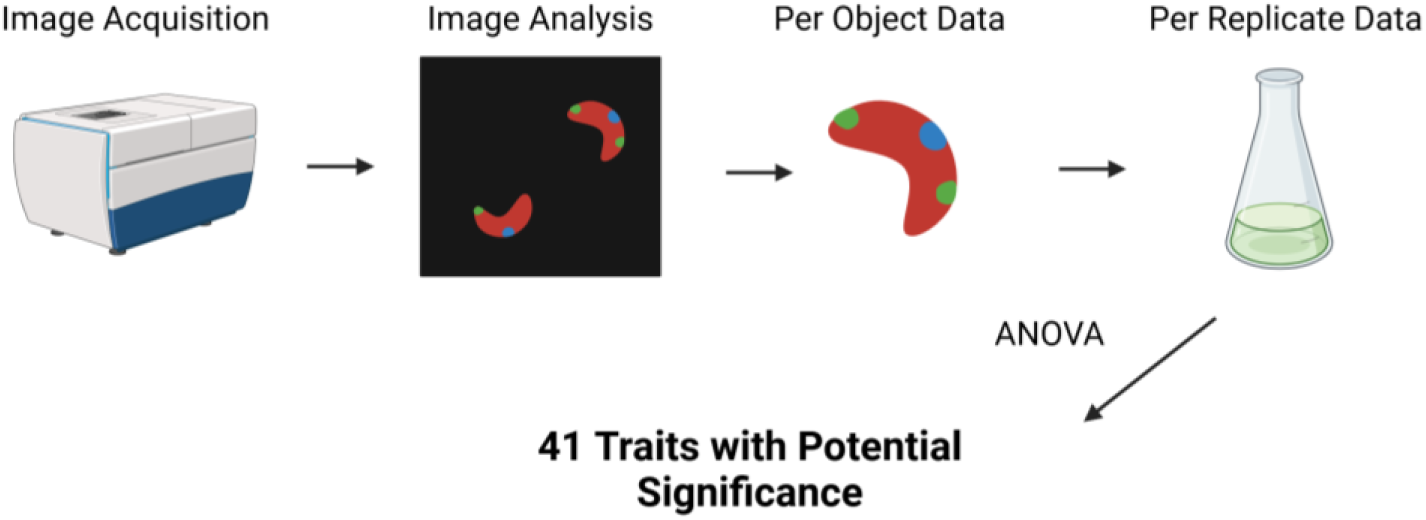
Diagram of the analysis carried out on imaging data, starting with data aggregation and ending with the use of ANOVA tests to identify traits with significant variation among treatments. Created in Biorender.

During the conduction of experiments, the original ImageXpress instrument being used was replaced due to technical issues. To account for any potential variation between images obtained on the old versus the new device, Z-score normalization was carried out before comparison of image data collected on different instruments. All imaging data was normalized to controls obtained on the same imaging system. Two controls were randomly selected from each week of exposure to include in this normalization and in subsequent image analysis. For statistical analysis of specific imaging data points (i.e. lipid droplet count) data was simply compared and visualized separately depending on instrument collected. Raw and normalized imaging data has been submitted as a public Mendeley Data dataset [26].

### 2.9 Statistical Analysis

Lipid droplet count, chlorophyll-a content, integrated intensity, and percent growth inhibition were analyzed separately from imaging data to determine if any varied significantly between treatments and controls. Growth data was not normally distributed, and was analyzed with a Kruskal-Wallis test and subsequent Dunn post-hoc test. Chlorophyll-a data was also not normally distributed, and was analyzed using a Kruskal-Wallis and Dunn test. Lipid droplet count and integrated intensity data were divided into two datasets for analysis: CP-CDs, CP-polymer, and nitrogen deprivation, and PEI-CDs, PEI polymer, and low light. They were then compared to the control. All analyses were visualized with ggplot2. All statistical analysis was carried out in R [27, 28].

Prior to statistical analysis, data with no variation or missing values was removed from the imaging analysis dataset. This resulted in a total of 152 morphological traits being used in further analysis. For analysis of morphological data, an ANOVA was used to screen the initial data set for variables with potential significant differences from the control (Figure 2). Variables deemed to have significant variance among treatments (p<0.05) were used for further analysis—41 variables were included in this dataset. A series of Tukey post-hoc tests were then run to find significant differences between treatments for each variable. Comparisons yielding a p-value of <0.05 were determined to be significant.

To determine if each CD treatment was most similar in its mechanism of action to nitrogen deprivation, growth in low light conditions, or the effects of its polymer, a random forest model was used. The model was trained on five known treatments: control, 10% nitrogen, CP 25 ug/L (polymer only), PEI 5 ug/L (polymer only), and low light. The known treatments were split into a training and test set, with 80% of samples in the training set and 20% in the test set. As the number of replicates varied among treatments, the upSample function was used to balance the dataset after splitting in R. A random forest model was then trained using the randomForest function in R [29]. The number of trees was set at 100, and number of variables sampled at each split was set as the square root of the number of variables used (12.288). The out-of-bag error rate of the model was determined to be 8.82%. For further validation, the previously designated “test” dataset was balanced and the model used to predict classes of each data point. The model correctly categorized the data 90% of the time. The model was then used to predict the class of each CD treatment.

To further visualize similarities and differences among treatments, a dendrogram was built. First, data was aggregated and mean values for each treatment were calculated. Then, the data was grouped and Euclidean distance was calculated using the dist function in R, resulting in a distance matrix. Hierarchical clustering was performed on the distance matrix using the hclust function and the ward.D2 method. Finally, the dendrogram was plotted in R.

## Results

### 3.1 Particle Characterization

Carbon dots were found to have similar size and charge as documented in previous studies. Box plots showing the results of particle characterization can be found in Figure S1. PEI-CDs were found to have a positive zeta potential of 24.05 ± 7.29 mV, reflecting their strong surface charge due to the presence of PEI functional groups, and a compact average diameter of 2.22 ± 0.21 nm, indicative of their uniform size. CP-CDs showed a negative zeta potential of –18.44 ± 10.10 mV, which can be attributed to the carboxyl functional groups on their surface, and a slightly larger mean size of 3.58 ± 0.30 nm, indicating the well-functionalized carboxyl groups on the surface of the CP-CDs. These results are consistent with previous studies using the same CDs [11].

### 3.2 Growth Inhibition

Relative to the control (mean 6.9389e-18 ± 3.0558% growth inhibition), the growth rate of *R. subcapitata* was significantly impacted during exposure to 10 ug/L of PEI-CDs (17.0717 ± 8.2617%), 10 ug/L of PEI polymer (14.9463 ± 5.0815%), and growth in reduced light conditions (17.6746 ± 2.4438%). Statistical analysis was performed using a Kruskal-Wallis and Dunn post-hoc test, where p-values less than 0.05 were considered significant. Although other treatments, especially growth in low nitrogen conditions, increased growth inhibition, these differences were not statistically significant (Figure 3).

**Figure 3.**
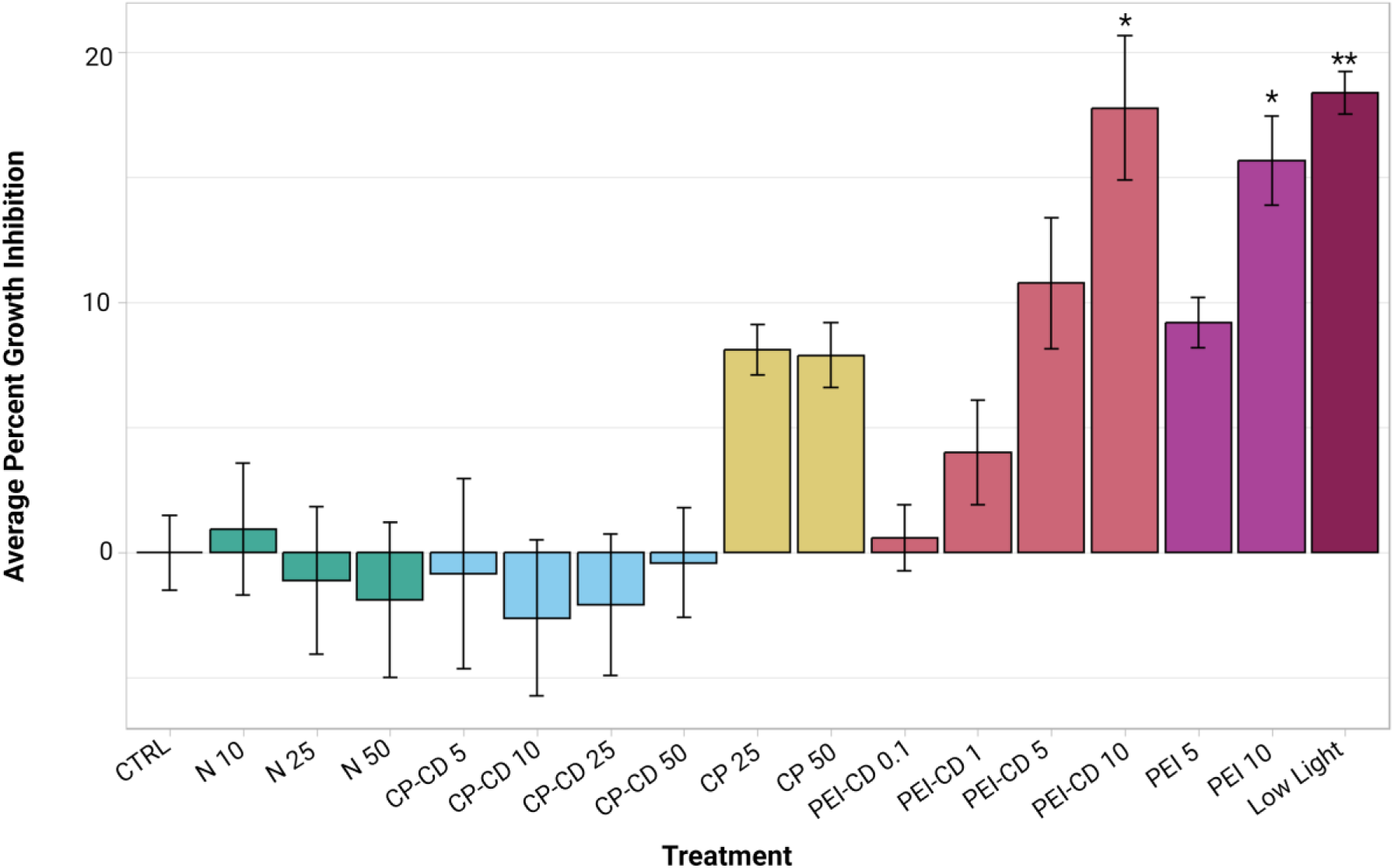
Average percent growth inhibition, plotted with ggplot2 [41]. Asterisks indicate significant difference from the control (p<0.05, Kruskal-Wallis and Dunn post hoc tests, n=8-12).

### 3.3 Chlorophyll-a Quantification

Results of chlorophyll-a quantification in PEI-CD, PEI polymer, and low light exposed cells are shown in Figure 4. Statistical analysis was performed using Kruskal-Wallis and Dunn post-hoc tests. When compared to the control (mean chlorophyll-a content of 8.7854e-7±6.8755e-8 ng/cell), only one treatment showed a significant difference in chlorophyll-a content: the low light treatment (mean chlorophyll-a content of 1.177e-6±1.2018e-7 ng/cell, p<0.05).

**Figure 4.**
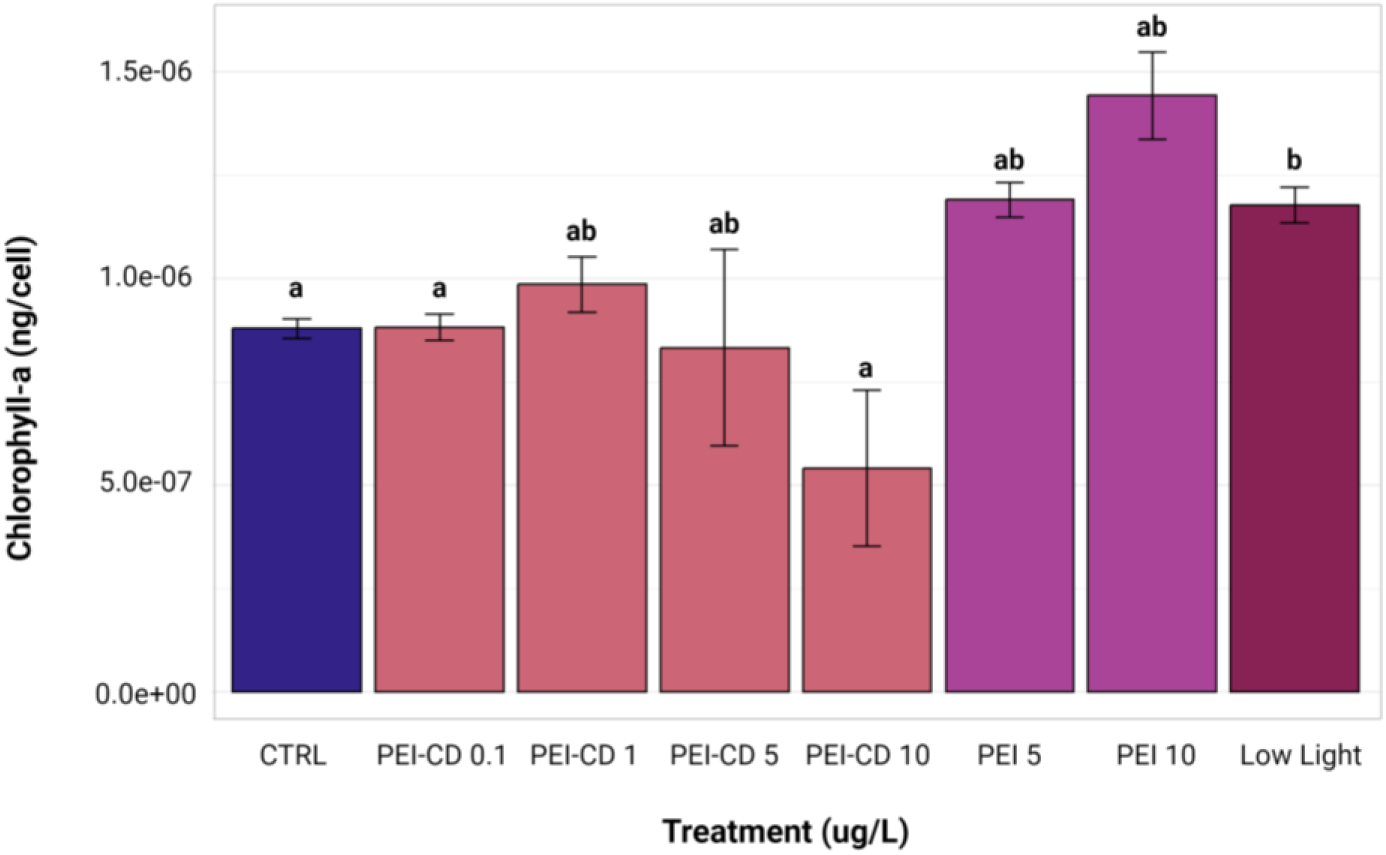
Chlorophyll-a content, corrected by cell count. Letters indicate significant differences among groups, with those having the same letter showing no significant difference from each other (p<0.05, Kruskal-Wallis and Dunn post hoc tests, n=8). Created with ggplot2 [41].

Chlorophyll-a content per cell was increased in this case. Although no other treatments were significantly different from the control, the PEI-CD 10 ug/L treatment (mean chlorophyll-a content of 5.4127e-7±5.3249e-7 ng/cell) did exhibit significantly lower chlorophyll-a content when compared to the low light treatment.

### 3.4 Lipid Droplet Count

Both PEI-CD 10 ug/L and low light treatments exhibited significant differences in lipid droplet count from the control (ANOVA and Dunnett’s Test post-hoc, p<0.05). Mean lipid droplet count per cell in control samples collected during PEI-CD exposures was 2.41±1.13. In contrast, cells exposed to PEI-CDs at a concentration of 10 ug/L had a mean lipid droplet count of 4.33±1.82, and those grown in low light conditions had a significantly lower LD count of 0.298±0.341 per cell (Figure 5). When low nitrogen, CP-CD, and CP polymer exposed cells were compared to their respective controls, no significant differences were observed.

**Figure 5.**
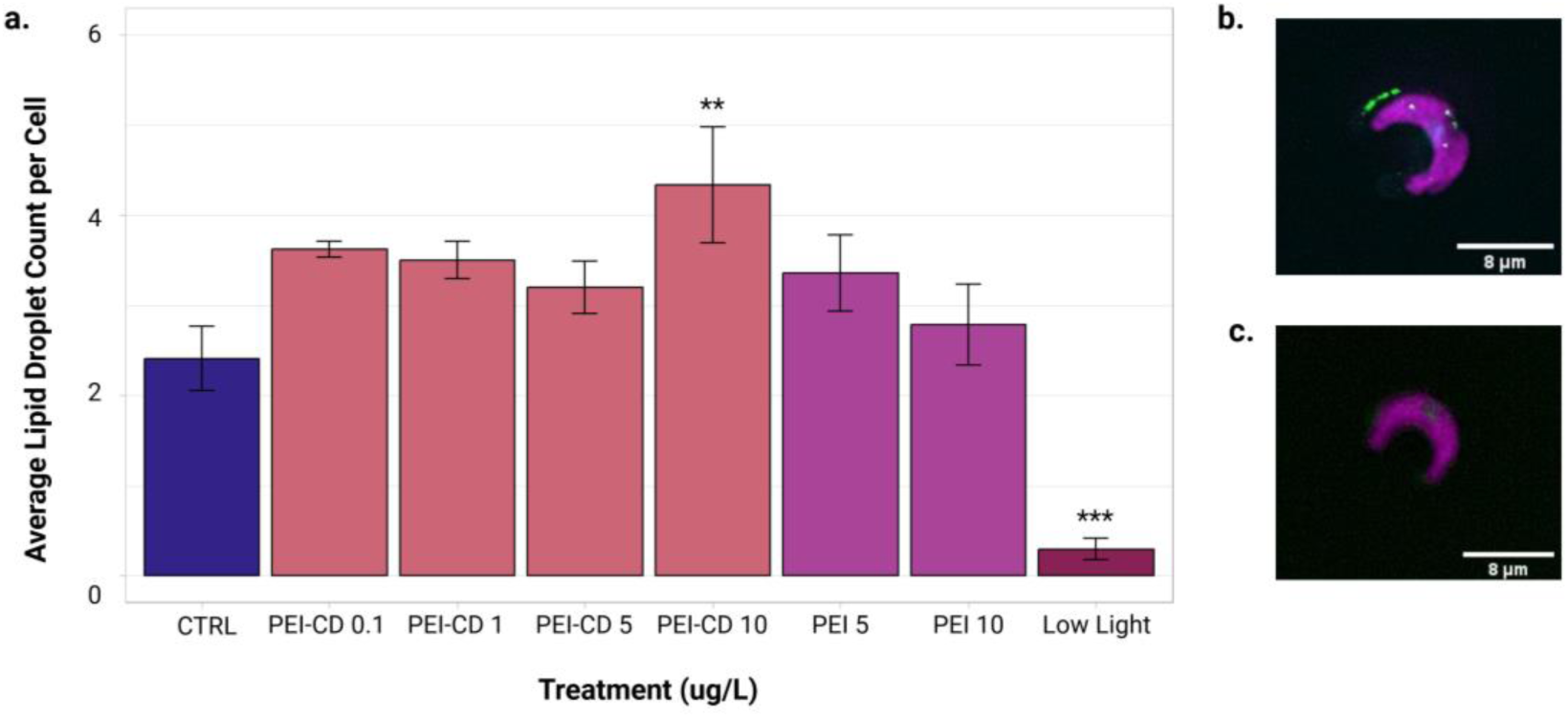
a. Mean lipid droplet count per cell of PEI and low light treatments, obtained from ImageXpress data analyzed in MetaXpress. Asterisks indicate significant difference from the control (ANOVA and Dunnett’s test post-hoc, p<0.05, n=8-10). Graph created with ggplot2 [41]. b. Image of a cell with lipid droplets present, edited in Fiji [40]. c. Example of cell without lipid droplets present, edited in Fiji [40].

### 3.5 Integrated Intensity

Three analyses of integrated intensity were performed. The first compared cells exposed to low nitrogen conditions or CP-CDs to their respective controls using a one-way ANOVA and subsequent Dunnett’s test. With a p-value of 0.0594 obtained from the one-way ANOVA, no significant differences were observed. CP polymer-only treatments were compared separately to their respective controls using a Kruskal-Wallis test with Dunn post-hoc. No significant differences were observed in this case (Kruskal-Wallis p = 0.742). Finally, PEI-CD, PEI polymer, and low light exposed cells were analyzed relative to their controls using Kruskal-Wallis and Dunn post-hoc tests. PEI-CD 10 ug/L (mean relative fluorescence 36654740.5812 ± 8765429.7110) and PEI polymer 10 ug/L (mean relative fluorescence 33181564.2400 ± 15410245.1351) both showed significantly decreased chloroplast integrated intensities relative to the control (mean relative fluorescence 48404984.8310 ± 5133518.8752) (Figure 6).

**Figure 6.**
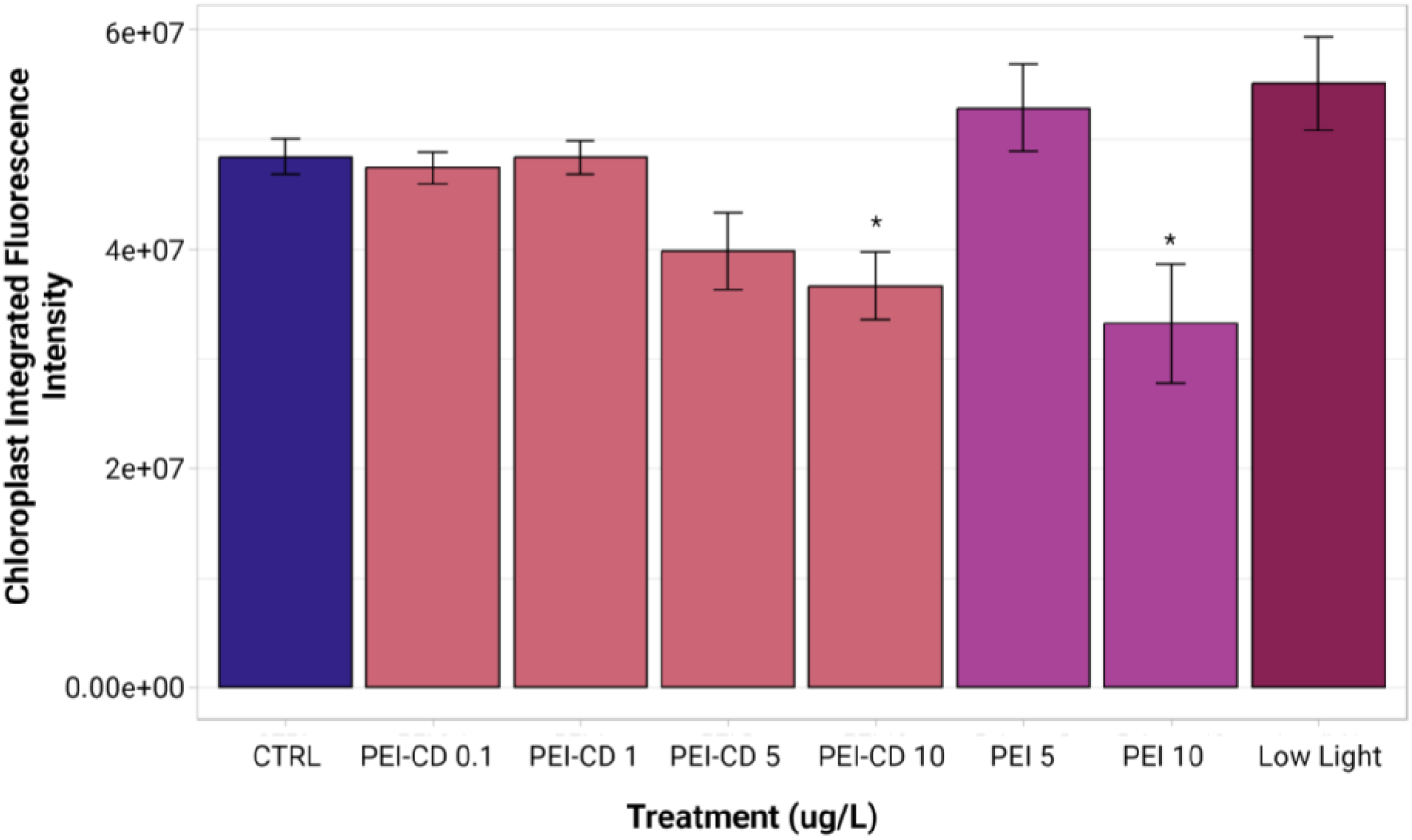
Mean integrated intensity of PEI and low light treatments. Asterisks indicate significant difference from the control (p<0.05, Kruskal-Wallis and Dunn post-hoc tests, n=8-10). Graph created with ggplot2 [41].

### 3.6 Morphological Characterization

Of the 152 morphological traits examined, 41 were determined to vary significantly amongst the treatments (ANOVA, p<0.05). To determine which traits varied significantly between specific treatments, a series of Tukey post-hoc tests were performed. A heatmap showing the results of these tests is available in Figure S2 (created in R the pheatmap package [30]. The two treatments with the most significantly different traits between them were low light and CP-CD 10 ug/L. To more clearly visualize the overall similarities and differences among treatments, a Euclidean distance dendrogram is shown in Figure 7. The dendrogram shows that PEI-CD treatments cluster together, but do not group with the low light or PEI polymer treatment. CP-CD treatments tended to cluster with low nitrogen treatments, suggesting similarities among these treatments.

**Figure 7.**
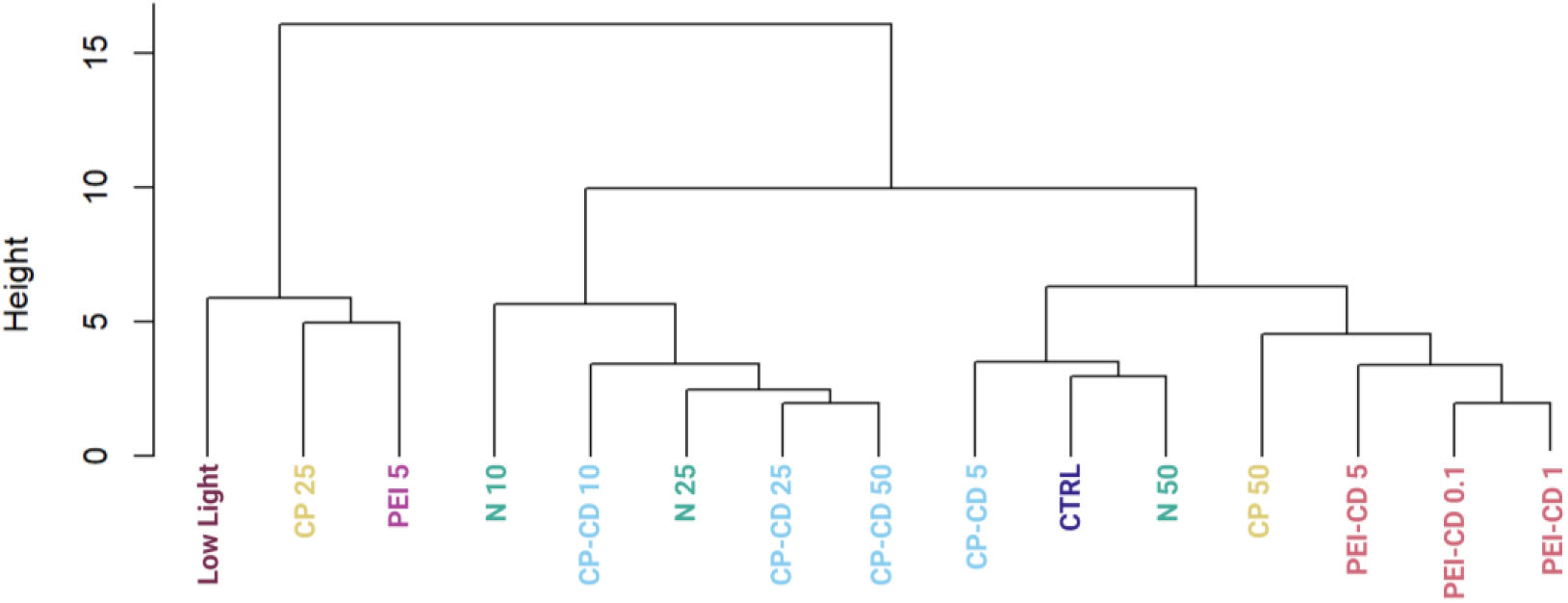
Dendrogram built using morphological data using Euclidean distance and hierarchical clustering.

### 3.7 Random Forest Categorization

A confusion matrix showing the results of random forest categorization of CD-exposed cells is shown in Figure 8. Cells exposed to PEI-CDs in doses of 0.1 and 1 ug/L were categorized as most similar to the control cells in 100% of cases. PEI 5 ug/L was categorized as control in 75% of cases, with 12.5% falling under the CP polymer 25 ug/L category and 12.5% in the PEI polymer 5 ug/L category. CP-CD exposed cells were most often categorized as the control or 10% nitrogen condition, with up to 62.5% of cases being categorized as 10% nitrogen depending on dose. However, CP-CD 5 ug/L was categorized as CP polymer 25 ug/L in 12.5% of cases. Overall, there was little similarity between the CD treatments and their respective polymer-only controls.

**Figure 8.**
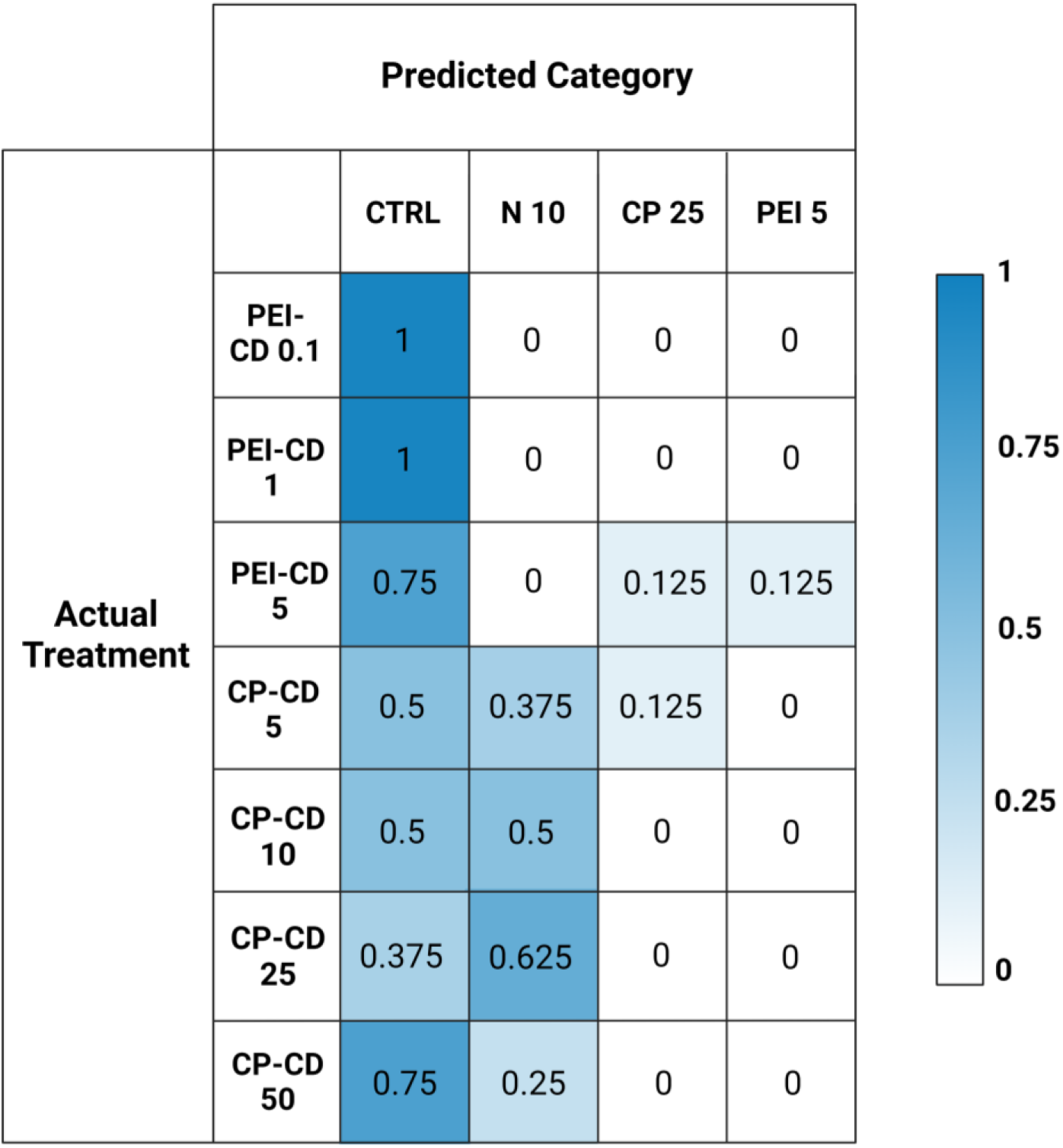
Matrix built from Random Forest classification data. Actual categories of classified data are on the left, with the predicted categories on the top. Numbers represent the proportion of samples assigned to that category.

## Discussion

Nanomaterials can increase lipid content in microalgae, but their ability to do so is clearly impacted by surface functionalization and charge. PEI-CDs reliably increase LD content, likely via direct interaction with the cell. This effect is nano-specific, though: morphological characterization strongly supports the conclusion that PEI-CDs act via a mechanism that is distinct from the PEI polymer alone. Specifically, while the PEI-CD 10 ug/L treatment increased LD content, PEI polymer at the same concentration did not. Further, when categorized by random forest, the PEI-CD 5 ug/L treatment categorizes as most similar to PEI polymer 5 ug/L only 12.5% of the time (Figure 8). A dendrogram built using a Euclidean distance matrix and hierarchical clustering groups PEI-CD treatments with each other, but not with the PEI polymer treatment included in the diagram (Figure 7). Rather, the PEI polymer 5 ug/L treatment clusters with a CP polymer treatment and the low light condition. The finding that PEI-CD induced LD production is nano-specific supports the idea that nanoparticles may have specific effects on microalgal lipid content and could serve as a tool in production of biofuels from microalgae.

Despite its lack of effect on LD content, data regarding the response of *R. subcapitata* to PEI polymer still provides valuable information—insight into the mechanism of action of PEI-CDs. Similarities observed between the impacts of PEI-CDs and PEI are important to note. Both significantly inhibited *R. subcapitata* growth and decreased chloroplast integrated intensity. A decrease in chloroplast integrated fluorescence intensity suggests either potential chloroplast damage and reduced photosynthetic performance, or CD quenching of chloroplast fluorescence pigments. This could be explained by the hypothesis that both PEI polymer, and the PEI polymer present on the CDs, damage the chloroplast membrane. PEI is known to damage membranes and increase membrane permeability [31, 32]. Furthermore, PEI-CDs are known to enter *R. subcapitata* cells [11]. It is possible, then, that both PEI and PEI-CDs enter microalgal cells, causing damage to intracellular membranes, such as that of the chloroplast. Although little work exists regarding the biological impacts of PEI on chloroplast function, such effects have been observed in mitochondria, where PEI has been shown to induce proton transport chain leakage [33, 34]. This data could explain the damage observed to the chloroplast in both PEI-CD and PEI polymer exposed cells.

The differences between PEI-CDs and PEI polymer may point to the causes of a nano-specific effect on lipid content. One difference between these treatments is observed in chlorophyll-a quantification data. While PEI-CD 10 ug/L is significantly different in chlorophyll-a content from the low light condition, PEI polymer 10 ug/L is not. This suggests a stronger shading effect, likely caused by greater aggregation, in cells treated with PEI polymer alone. This aligns with lipid content data. PEI polymer does not induce increased lipid production, while PEI-CDs do. Growth in low light conditions significantly decreased lipid content. PEI polymer alone is clearly more similar to a shading effect.

Taking both the similarities and differences between PEI-CD and PEI polymer response into account, one can hypothesize that uptake is key to PEI-CD induced lipid production. PEI-CDs are known to be taken up by *R. subcapitata*, and evidence in this study suggests the uptake is to a higher extent than PEI polymer. PEI polymer is more likely to remain on the cell surface, interacting electrostatically with microalgal cells to form aggregates. These aggregates reduce light availability to the cell, resulting in a compensatory chlorophyll-a content increase. However, some PEI polymer still enters the cell, causing some chloroplast damage and decreased chloroplast integrated intensity.

Along with emphasizing the importance of uptake, the findings of this study also exemplify the charge dependent mechanisms at play in *R. subcapitata*’s response to carbon dot exposure. CP-CDs have been documented to be taken up to a lesser extent than PEI-CDs [11]. While previous evidence has shown potential for CP-CDs to induce increased LD content, this result is more variable than PEI-CD induced LD production. In the current study, CP-CD treatment did increase LD content, but not significantly. The CP-CD treatment increasing LD content the most was 25 ug/L, which increased LD content to 0.8308 ± 0.3040 mean lipid droplets per cell, compared to 0.5522 ± 0.3221 per cell in the control (p =0.4, Welch’s One-Way ANOVA and Games-Howell Multiple Comparison tests). No differences between the carbon dot and polymer treatment were observed, as neither showed statistical differences from the control. Evidence was found to suggest that CP-CDs interact with ammonium in OECD media, though. This indirect interaction could explain previously observed impacts to *R. subcapitata* lipid content.

Morphological data is the primary evidence for this hypothesis. Random forest classification shows that CP-CD exposed cells group with nitrogen deprivation (10% nitrogen relative to control) 37.5% of the time for CP-CD 5 ug/L, 50% for 10 ug/L, 62.5% for 25 ug/L, and 25% for 50 ug/L. In a dendrogram built by calculating Euclidean distance and completing hierarchical clustering on the morphological data of all treatments, CP-CD 10, 25, and 50 ug/L cluster with 10% and 25% N. CP-CD 5 ug/L clusters with 50% N and the control. This further suggests a similarity between the CP-CD and nitrogen deprivation treatments.

Positively charged CDs clearly have the potential to induce increased lipid droplet production in microalgae at low doses (10 ug/L). But the results of this study leave the effectiveness of CD use for biofuel production in question. More work is required to answer this question completely.

However, a few findings may provide insight into how CDs should be studied in the field of algal-based biofuels. This is a much lower concentration than those used in other studies wherein nanoparticle exposure positively impacted LD content. For example, a 2014 study by Sarma et al utilized 1 g/L of MgSO_4_ nanoparticles to induce a percent increase in lipid content of 97.67 ± 6.9% [35]. While the low dose needed to see significant changes in lipid content with PEI-CDs is promising, these particles also induce growth inhibition at these low doses. Many factors besides charge influence nanoparticle toxicity, though, and modifying these could yield a nanoparticle with similar impacts on LD content but lowered toxicity. One factor impacting toxicity is NP size. Smaller nanoparticles typically cause higher rates of toxicity [36, 37]. Using a larger nanoparticle could decrease toxicity—however, it could also decrease uptake. Nanoparticles too large to pass through algal cell wall pores, which often range in size between 5 and 20 nm, are unlikely to be taken up [38]. As the PEI-CDs used in this study were found to be taken up by *R. subcapitata*, and fell within the size range thought to be taken up by microalgae (PEI-CDs: 2.22 ± 0.21 nm; CP-CDs: 3.58 ± 0.30 nm) it is very possible that increasing particle size will reduce uptake and change the cellular response to the CDs.

Perhaps a more promising modification to reduce toxicity while maintaining impacts to lipid production is changing particle surface charge density [39]. The reduction of surface charge density has been shown to decrease nanoparticle toxicity in mammalian cells [40] and reduce interactions with plant cell walls [39]. Using a PEI-CD with lower surface charge density could lower toxicity, but the impacts of surface charge density on microalgal lipid content are unknown and should be further investigated. A particle with lowered surface charge density would likely exhibit decreased toxicity and could induce similar increases to lipid content. In summary, PEI-CDs show promise in their ability to increase microalgal lipid content. Simple nanoengineering strategies could be applied to reduce their toxicity and improve lipid yield when they are added.

In contrast, the evidence in this study suggests that CP-CDs may not impact lipid content differently from nitrogen deprivation, at least at concentrations at or below 50 ug/L. If this is the case, the use of CP-CDs would not be more economically or environmentally favorable than nitrogen deprivation alone. Additionally, no evidence was found to suggest that CP-CDs impact *R. subcapitata* differently than CP polymer alone. The lack of nano-specific effects suggests that CP-CDs may not be the most economical method for impacting microalgal LD content.

### Conclusion

This study is one of the first to demonstrate that nanoparticles can have nano-specific effects on microalgal lipid production. The use of PEI-CDs increases lipid content in *R. subcapitata*, while the PEI polymer alone did not. PEI-CDs are not a perfect tool for increasing TAG content in microalgae—low levels of growth inhibition occurred in conjunction with increases in LD count. However, nanoengineering strategies, such as modifying surface charge density, could be applied to these nanoparticles in the future. Additionally, data regarding the mechanism of action of PEI-CDs was obtained. This data suggests that shading, a previously hypothesized mechanism of action of PEI-CDs, is likely not the primary cause of PEI-CD toxicity. Uptake, and subsequent cellular damage, is much more likely.

Contrastingly, CP-CDs were found to share a likely mechanism of action with nitrogen deprivation, and did not exhibit nano-specific effects. Any previously observed effects of CP-CDs on TAG content could be related to this mechanism. Therefore, CP-CDs are a less useful tool in the production of biofuels from microalgae. Similar effects can be obtained by simply reducing nitrogen availability to microalgae.

This study has significant implications for those studying the potential role nanotechnology could play in biofuel production from algae. Findings suggest that positively charged nanoparticles can have nano-specific effects on TAG content at relatively low (10 ug/L) concentrations. It also shows that the composition of the nanoparticle core does not determine the impacts of the particle on LD content when functionalization is used. Surface functionalization and charge are critical factors in the impact of nanoparticles on TAG content and mediate the mechanisms of action by which these particles act. Future work should further investigate the use of positively charged nanoparticles with different charge densities on algal biofuel production, aiming to reduce the toxicity of these particles in more realistic conditions such as algal bioreactors so that they can serve as an effective tool in sustainable fuel production.

## Acknowledgements

This material is based upon work supported by the National Science Foundation under Grant No. CHE-2001611, the NSF Center for Sustainable Nanotechnology. The CSN is part of the Centers for Chemical Innovation Program. Figures created with Biorender.

## CRediT Author Statement

**Emma McKeel**: Conceptualization, Investigaton, Formal Analysis, Writing-Original Draft **Hye-In Kim**: Investigation, Writing-Review and Editing **Su-Ji Jeon**: Investigation, Writing-Review and Editing **Britta McKinnon**: Investigation, Writing-Review and Editing **Juan Pablo Giraldo**: Supervision, Funding Acquisition, Writing-Review and Editing **Rebecca Klaper**: Supervision, Funding Acquisition, Writing-Review and Editing

