## Supplemental Information for "Mechanism of Nanomaterial-Induced Lipid Droplet Formation in *Raphidocelis subcapitata* is Mediated by Charge Properties"

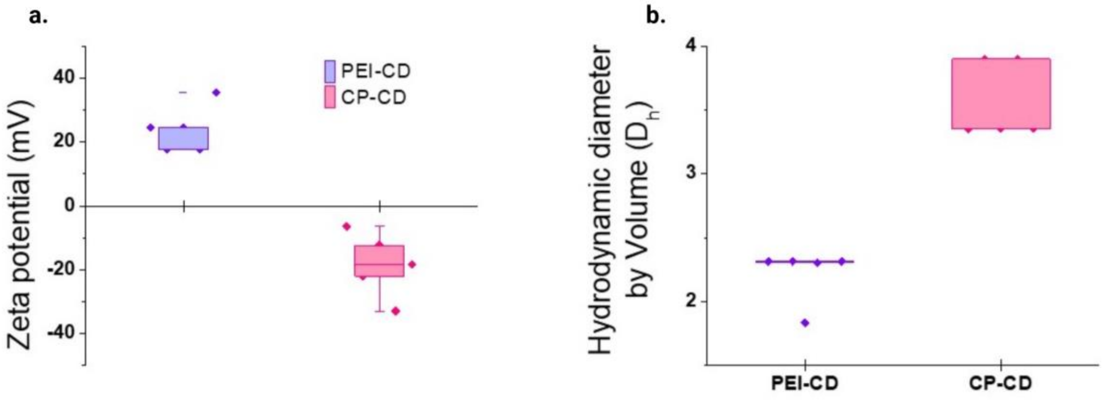


**Figure S1**. Box plots of DLS data obtained for PEI and CP-CDs: 1.a. shows zeta potential data, while 1.b. displays hydrodynamic diameter.

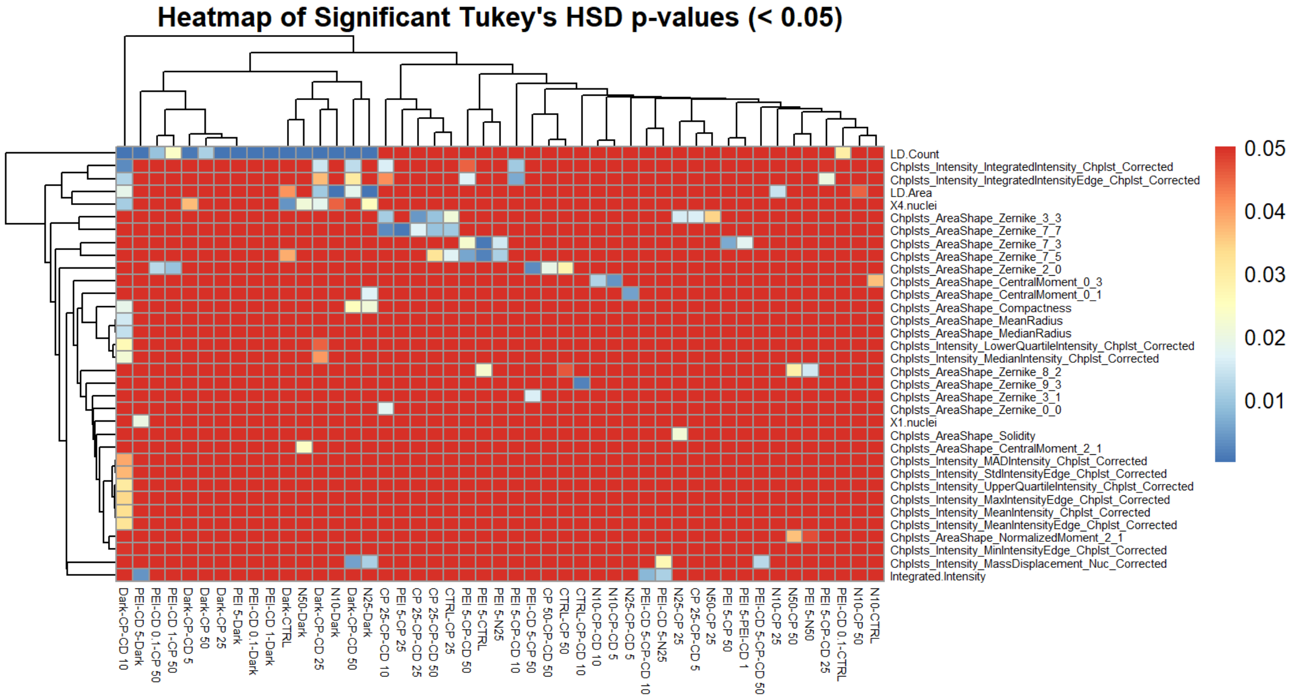


**Figure S2**. Heatmap of every treatment comparison with traits varying significantly between them. P-values greater than or equal to 0.05 are red, while other colors represent a p-value less than 0.05.

**MetaXpress Analysis Pipeline**

Image analysis in MetaXpress was carried out as described by McKeel et al:

These settings were used at 60x magnification and may need to be adjusted for other objectives. All modules are located under the “Segment” tab.

Step 1: Set image names and select Cy5, DAPI, and GFP channels.

Step 2: Filter DAPI image using Top Hat function. Set pixel size to 16, filter shape to circle, and name filter output image.

Step 3: Find nuclei using “Find Round Objects” function. Choose Top Hat filtered DAPI image as source. Set approximate minimum width to 1 μm and approximate maximum width to 2.11 μm. Set intensity above local background to 700, and name result “nuclei.”

Step 4: Using result “Nuclei” from step 3, mark the center of each nucleus as a single pixel using “Mark Object Centers” function. Name the result. This will allow for nuclei per cell counts, as number of pixels per cell will correspond to the nuclei count per cell.

Step 5: Create an image from the Mark Object Centers mask created in step 4 using the “Mask to Image” function. The mask to image here was named “Nuclei center Mask to Image.”

Step 6: Use “Top Hat” function to filter Cy5 images. Choose Cy5 as source, pixel size 22, and filter shape circle. Name result (named “TopHat Cell” here).

Step 7: Use the “Gaussian Filter” function to blur “TopHat Cell,” which aids in recognition of the sickle-shaped *R. subcapitata* cell. Rename (“blurred cell” in this example).

Step 8: Use function “Adaptive Threshold” to identify cells. Set source as “blurred cell,” approximate minimum width as 2.3 μm and maximum width as 4 μm. Use 1000 as intensity above local background and name result (“Cell” used as name here).

Step 9: Create a filter mask from “Cell” result using the “Filter Mask” function. Choose Integrated Intensity as as “Measurement,” “Range Filter” for “Filter Type” and set both minimum and maximum values to 1. Name “Filter Mask 1.” This identifies all cells with 1 nuclei.

Step 10: Use the “Filter Mask” function again. Choose “Nuclei center Mask to Image” as the image source and “Cell” as the mask source. Use “Integrated Intensity as measurement, “RangeFilter” as filter type, and set both minimum and maximum values to 2. Name this output “Filter Mask 2.” This identifies all cells with 2 nuclei.

Step 11: Use the “Filter Mask” function again with same settings as steps 9 and 10. This time, change the minimum and maximum values to 4. This identifies all cells with 4 nuclei. Name “Filter Mask 4.”

Step 12: Create a final “Filter Mask” as above, but change minimum value to 5, and do not include a maximum value. This creates a filter for any cells which have more than 4 nuclei.

Step 13: Use “Top Hat” function to filter GFP images. Set source as “GFP,” size to 10 pixels, and filter shape to “Circle.” Name “Top hat gfp.”

Step 14: Use a Gaussian filter to filter the Top Hat GFP image.

Step 15: Use “Find Round Objects” function to identify lipid droplets. Set source as “Top hat gfp,” approximate minimum width to 0.8 µm, approximate maximum width to 2.55 µm, and intensity above local background to 1200. Name result “lipid droplets.”
